# Neutralizing antibody-independent immunity to SARS-CoV-2 in hamsters and hACE-2 transgenic mice immunized with a RBD/Nucleocapsid fusion protein

**DOI:** 10.1101/2021.09.16.460663

**Authors:** Julia T. Castro, Marcílio J. Fumagalli, Natalia S. Hojo-Souza, Patrick Azevedo, Natalia Salazar, Bruna Rattis, Simone G. Ramos, Lídia Faustino, Gregório G. Almeida, Livia I. Oliveira, Tomas G. Marçal, Marconi Augusto, Rubens Magalhães, Bruno Cassaro, Gabriela Burle, Daniel Doro, Jorge Kalil, Edson Durigon, Andrés Salazar, Otávia Caballero, Alexandre Machado, João S. Silva, Flávio da Fonseca, Ana Paula Fernandes, Santuza R. Teixeira, Ricardo T. Gazzinelli

## Abstract

The nucleocapsid (N) and the receptor binding domain (RBD) of the Spike (S) proteins elicit robust antibody and T cell responses either in vaccinated or COVID-19 convalescent individuals. We generated a chimeric protein that comprises the sequences of RBD from S and N antigens (SpiN). SpiN was highly immunogenic and elicited a strong IFNγ response from T cells and high levels of antibodies to the inactivated virus, but no neutralizing antibodies. Importantly, hamsters and the human Angiotensin Convertase Enzyme-2-transgenic mice immunized with SpiN were highly resistant to challenge with the wild type SARS-CoV-2, as indicated by viral load, clinical outcome, lung inflammation and lethality. Thus, the N protein should be considered to induce T-cell-based immunity to improve SARS-CoV-2 vaccines, and eventually to circumvent the immune scape by variants.

---

The role of T cells on protective immunity to SARS-CoV-2 is poorly understood. *In vitro*, neutralizing antibodies (nAbs) bind to the Receptor Binding Domain (RBD) from Spike (S) protein and prevent the interaction of SARS-CoV-2 with the Angiotensin Convertase Enzyme-2 (ACE-2) and posterior host cell invasion ^1^. While the levels of neutralizing antibodies (nAb) elicited by vaccination or infection are taken as the main predictors of protective immunity, a causal-effect relationship remains to be established ^2-4^. Here, we report that immunized human-ACE-2 transgenic mice (K18-ACE-2) as well as hamsters, develop strong protective immunity to SARS-CoV-2 infection, even in the absence of nAbs. Hamsters and K18-ACE-2 mice immunized with a nucleocapsid (N) and RBD chimeric protein (SpiN), with no RBD conformational B cell epitopes, presented high levels of circulating anti-viral specific antibodies and T cells. Despite the high levels of antibodies specific for the N and RBD proteins, we were unable to detect nAbs in the sera of either rodent species vaccinated with SpiN. Regardless, immunization with SpiN induced a robust immune-mediated protection to experimental challenge with the wild type (Wuhan strain) SARS-CoV-2, which was otherwise lethal to non-vaccinated mice. These unexpected findings of nAb-independent protective immunity elicited by N fused to unfolded RBD protein might be of value in improving current COVID-19 vaccines.

Since the end of 2019, over 200 million cases and four million deaths from COVID-19 have been reported worldwide. Except for the use of inactivated SARS-CoV-2, all other COVID-19 vaccines that are currently in use or in advanced stages of development, are based on the S protein ^5,6^. However, positive selection of variants with non-synonymous mutations in the Spike gene has been observed ^7,8^. These SARS-CoV-2 variants have conformational changes in the RBD region of the S protein, augmenting their affinity to ACE-2 as well as the ability to escape from nAbs ^9^. While maintaining fitness, these conformational changes in the Spike protein are associated with enhanced viral infectivity and spread in humans ^10,11^. Indeed, the efficacy of the current vaccines targeting conformational RBD epitopes has been challenged by the emergence of new variants ^10,12-14^.

Evidence of the importance of T cells in mediating immunity to SARS-CoV-2 have accumulated over the last year ^15^. A coordinated response of CD4^+^ T cells, CD8^+^ T cells and nAbs seems to be ideal for a favorable outcome of disease ^16^. Importantly, asymptomatic patients have low, often undetectable, levels of anti-SARS-CoV-2 antibodies and nAbs. In contrast, patients that develop moderate or severe disease present intermediate and high levels of circulating nAbs ^17-24^. Furthermore, multiple T cell epitopes have been identified in SARS-CoV-2 proteins, some of which present homology to polypeptides from other coronavirus that circulate in the human populations that might explain the resistance of some seronegative individuals to symptomatic COVID-19 ^4,25,26^. Altogether, these studies suggest an important role of effector T cells in mediating resistance to primary infection with SARS-CoV-2.

With the aim of developing a vaccine that induces a strong T cell mediated immunity, we considered the N protein together with the canonical S antigen that is employed in most COVID-19 recombinant vaccines. The N protein is the most abundant SARS-CoV-2 protein expressed in the host cell cytosol ^27^. It is thus likely to be presented for cytotoxic CD8^+^ T cells via HLA-I through the endogenous pathway. The inclusion of the RBD region shall contribute with helper T cells for B lymphocytes and antibody response to the main SARS-CoV-2 antigens, including the nAbs. In addition, both N and S proteins have been shown highly immunogenic for CD4^+^ T and CD8^+^ T cells and B lymphocytes ^4,16,26,28,29^.

First, we performed *in silico* analysis of the N and S proteins to identify the regions that are enriched on T cell epitopes ^30,31^ (**Extended Data Fig. 1a-d, Fig. 1a**). We found that in the S protein, the RBD segment has the greatest prevalence of potential T cell epitopes. It is also shown in **Fig. 1a** that the N protein is highly enriched on both CD4^+^ T and CD8^+^ T cell epitopes. Next, we analyzed the N and S protein sequences of 63 and 61 isolates, respectively, categorized as the three variants of concern (VOC; Alpha-B.1.1.7, Beta-B.1.351, Gama-P.1), and two variants under monitoring (Zeta-P.2, Epsilon-B.1.427/B.1.429) of SARS-CoV-2 distributed worldwide (**Extended Data Fig. 2a**,**b**) ^32,33^. The peaks with circles indicate the position of most frequent amino acid changes in the N and S proteins, and the height of the peaks indicates the frequency that these changes occur. The black and red lines below the bars representing the N and S proteins indicate each of the putative CD8^+^ T and CD4^+^ T cell epitopes, respectively (**Fig. 1a**). Importantly, few of these epitopes overlapped with sites of amino acid mutations that are associated with the variants (peaks with purple circles). The vast majority of the putative T cell peptides from N protein and RBD domain were conserved (**Fig. 1a**).

**Figure 1:**
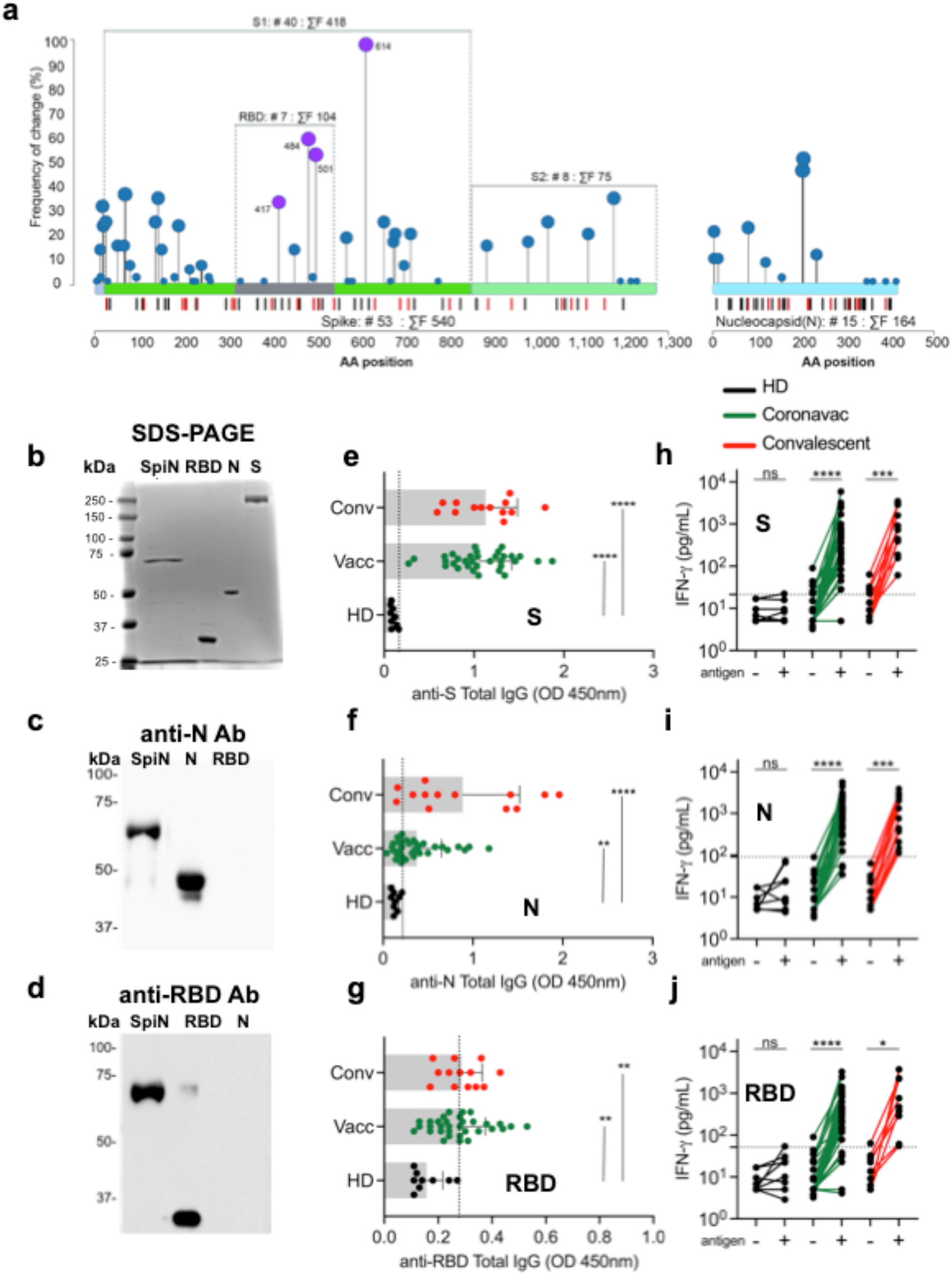
Human antibody and IFNγ responses to N and RBD polypetides. **a**, Needle plot, indicating the number of amino acid divergence points in the protein sequence of Nucleocapsid (N) (63 isolates) and Spike (S) (61 isolates) in relation to the N and S sequences from the SARS-CoV-2 lineage B (Wuhan). The peaks with circles indicate the position of most frequent amino acid changes. The height of the peaks indicates the frequency of the changes in each divergent point. The blue and purple circles indicate common mutations and those observed on variants of concern, respectively. The sum of the amino acid changes for each segment (S1, RBD and S2) of the S and N proteins is also shown. The vertical black and red lines below the bars illustrating the N and S polypeptides indicate each of the putative CD4^+^ T and CD8^+^ T cell epitopes identified by *in silico* epitope prediction. **b**, SDS-PAGE of purified SpiN, RBD, N and S proteins. **c, d**, Western blots of purified SpiN, RBD and N proteins using rabbit polyclonal anti-N and a mouse monoclonal anti-RBD antibodies are shown in panels. **e-g**, The levels of IgG antibodies specific for S, N and RBD proteins found in sera from healthy controls (HC), vaccinated and convalescents, as indicated. **h-j**, The IFNγ response, respectively, to the S, N and RBD proteins by the PBMCs from HC, vaccinated and convalescents. The lines link the levels of IFNγ produced by PBMCs cultured in the absence and presence of antigens, as indicated. IgG antibodies measurements were analyzed through Two-way ANOVA followed by Dunn’s multiple comparisons test. Statistical analysis of IFNγ production was performed using Wilcoxon-matched pairs signed rank. NS indicate that difference is not statistically significant. ** P < 0.01 and **** P < 0.0001.

Although no data are available, the N protein is considered a candidate for a COVID-19 vaccine ^6,34,35^. Since the N protein is highly enriched for putative T cell epitopes, we used this structural protein as the basis of our vaccine. Importantly, N protein is the most abundant viral protein in infected cells, being highly expressed in their cytosol ^27^, and thus readily available for processing and presentation to cytotoxic T cells via HLA-I. In contrast, the S protein is directed to the host cell membrane. In addition, we included in our chimeric protein the RBD region from the S antigen, the second most abundant SARS-CoV-2 protein in the host cells. The RBD is also highly immunogenic for helper T cells ^16,26,27^ that upon viral infection, could rapidly promote the production of anti-S nAbs. We hypothesize that a vaccine containing multiple linear T cell epitopes would be effective against SARS-CoV-2 variants that evade the protective nAbs based on a restricted number of nonsynonymous mutations in their genome ^8,10,11,13,14^.

Polyacrylamide gels (**Fig. 1b**) and Western blots (**Fig. 1c,d**) show the highly purified N (∼45 kDa) and RBD (25∼ kDa) proteins expressed in *Escherichia coli* as well as S surface antigen (∼98 kDa) obtained from eukaryotic cells. The SpiN chimeric protein that contains the N protein and the RBD region from S protein has an apparent molecular weight of 70 kDa (**Fig. 1b-d**). The proteins were purified either on a nickel column (RBD and N) or by ion exchange chromatography (SpiN). The immunoblots were generated with either rabbit polyclonal anti-N (**Fig. 1c**) or anti-RBD (**Fig. 1d**) sera and show that SpiN protein is recognized by both antibodies. To ensure that N or RBD proteins are recognized, both by antibodies and T cells from humans, we used samples from COVID-19 convalescent and individuals that have been vaccinated with an inactivated virus vaccine (CoronaVac). The levels of circulating antibodies in sera from convalescents were variable, whereas from vaccinated individuals were more uniform. This is likely to be due to the time of serum samples collection, which varied from 2 to 8 months after viral detection by RT-PCR in convalescents and 1 to 2 months after the second dose in vaccinated individuals. Convalescent individuals developed a high, but variable antibody response to S (**Fig. 1e**) and N proteins (**Fig. 1f**) and low response to the unfolded RBD expressed in bacteria (**Fig. 1g**). Interestingly, vaccinated individuals showed a low antibody response to both N and RBD proteins, while the response to N protein was higher in most convalescent individuals (**Fig. 1f**). Importantly, S (**Fig. 1h**), N (**Fig. 1i**) and RBD (**Fig. 1j**) proteins induced a strong T cell response in most patients, consistent with the high content of putative T cell epitopes. The vaccinated individuals, but not seronegative healthy donors (HD), also showed a robust IFNγ response to the N and RBD recombinant proteins.

Next, we evaluated the immunogenicity and whether immunization with RBD, N and SpiN recombinant proteins protects mice against SARS-CoV-2 challenge. Mice were immunized with either recombinant N, RBD or SpiN by giving two intramuscular doses scheduled 21 days apart (**Fig. 2a**). As immunological adjuvants we used the synthetic polyinosinic-polycytidylic acid (Poly I:C) mixed with the stabilizers carboxymethylcellulose and polylysine (Poly-ICLC, Hiltonol) or a MF59-like squalene-based oil-in-water nano-emulsion (Addavax, Vac-Adx-10 InvivoGen). We have chosen these adjuvants because they were shown to induce an effective immunity to influenza that also infects humans through the respiratory tract ^36-38^. The Poly-IC derivatives are potent activator of Toll-Like Receptor 3 (TLR3) and other cytosolic innate immune receptors that recognize viral double stranded RNA and favors T cell-mediated immunity. It has also been used in multiple clinical trials for cancer therapy ^39-41^. The MF59-like squalene-based oil-in-water nano-emulsion formulation lacks agonists for innate sensors, but is used in a commercially available vaccine and shows a high performance ^38^.

**Figure 2:**
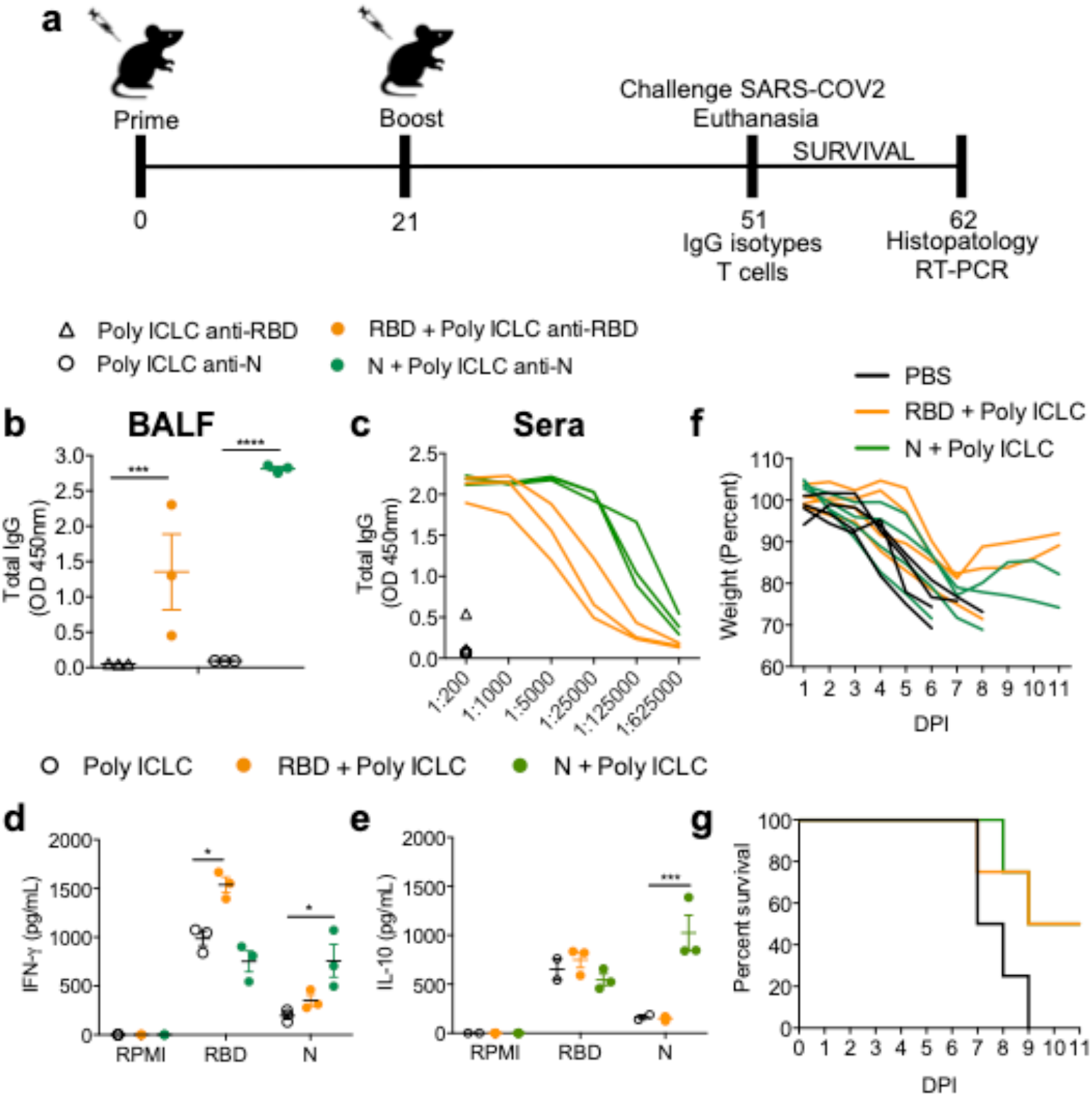
Evaluation of immune response and protection elicited by vaccination with RBD or N proteins associated with Poly ICLC. **a**, The immunization protocol used in the experiments shown in **Figures 2 to 6** to analyze the immune response as well as protection against SARS-CoV-2 infection. **b**, Antigen-specific IgG antibodies measured in the bronchoalveolar lavage (BALF) at 1:1 and serially diluted sera from immunized mice **(c)**. The levels of IFNγ **(d)** and IL-10 **(e)** were measured on culture supernatant of splenocytes stimulated with RBD or N antigens. **f, g**, Body weight and survival of mice immunized with either RBD or N associated with Poly ICLC and challenged with the Wuhan strain of SARS-CoV-2. **b-g**, Data are representative of two independent experiments. **b-e**, n=3 mice/group. **f**,**g**, n=4 mice/group. Statistical analysis of IgG measured in BALF was performed using two-tailed unpaired t-test, t=86.39 df=4. Cytokine measurements were analyzed through Two-way ANOVA followed by Tukey’s multiple test, df=30 for IFNγ; df=21 for IL-10. * P < 0.05, *** P < 0.001 and **** P < 0.0001.

The results demonstrate that either recombinant RBD or N protein associated to Poly ICLC are highly immunogenic inducing, respectively, high levels of anti-RBD or anti-N antibodies, both in the bronchioalveolar fluid (BALF) (**Fig. 2b**) and sera (**Fig. 2c**) of vaccinated mice. We also observed a strong IFNγ response (**Fig. 2d**), and less so for IL-10 (**Fig. 2e**), in splenocytes stimulated with these recombinant proteins. The K18-ACE-2 mice are a model of severe disease ^42^, and were used to evaluate the efficacy of immunization with either N or RBD recombinant proteins associated to Poly ICLC. Our results show that immunization with either protein resulted only in partial protection to SARS-CoV-2 challenge, as indicated by body weight loss and mortality (**Fig. 2f, g**).

Importantly, we report that immunization with adjuvanted SpiN fusion protein induced a robust viral-specific T cell and antibody responses, which are highly efficacious in protecting against experimental challenge with the SARS-CoV-2. Sera from mice immunized with SpiN associated to Poly ICLC showed very high titers of IgG antibodies specific for RBD (1:5,000) and N proteins (1:25,000) (**Fig. 3a**) as well as inactivated virus (1:5,000), but low titers of anti-S antibodies (**Fig. 3b**). The levels of anti-RBD and anti-N antibodies were equally higher in mice immunized with SpiN adjuvanted with Addavax **(Extended Data Fig. 3a)**. The levels of IgG anti-N (1:400) **(Fig. 3c)** as well as anti-SARS-CoV-2 (1:200) (**Fig. 3d**) were higher in sera from COVID-19 convalescent individuals than from healthy controls, but relatively low when compared to immunized mice. In contrast to immunized mice, the titers of anti-RBD were low (**Fig. 3c**) and anti-S (1:200) high (**Fig. 3d**) in sera of convalescent individuals. The results presented in **Figures 3e and 3f** show the increased levels of antibodies anti-N and anti-RBD in the BALF of mice vaccinated with SpiN, respectively.

**Figure 3:**
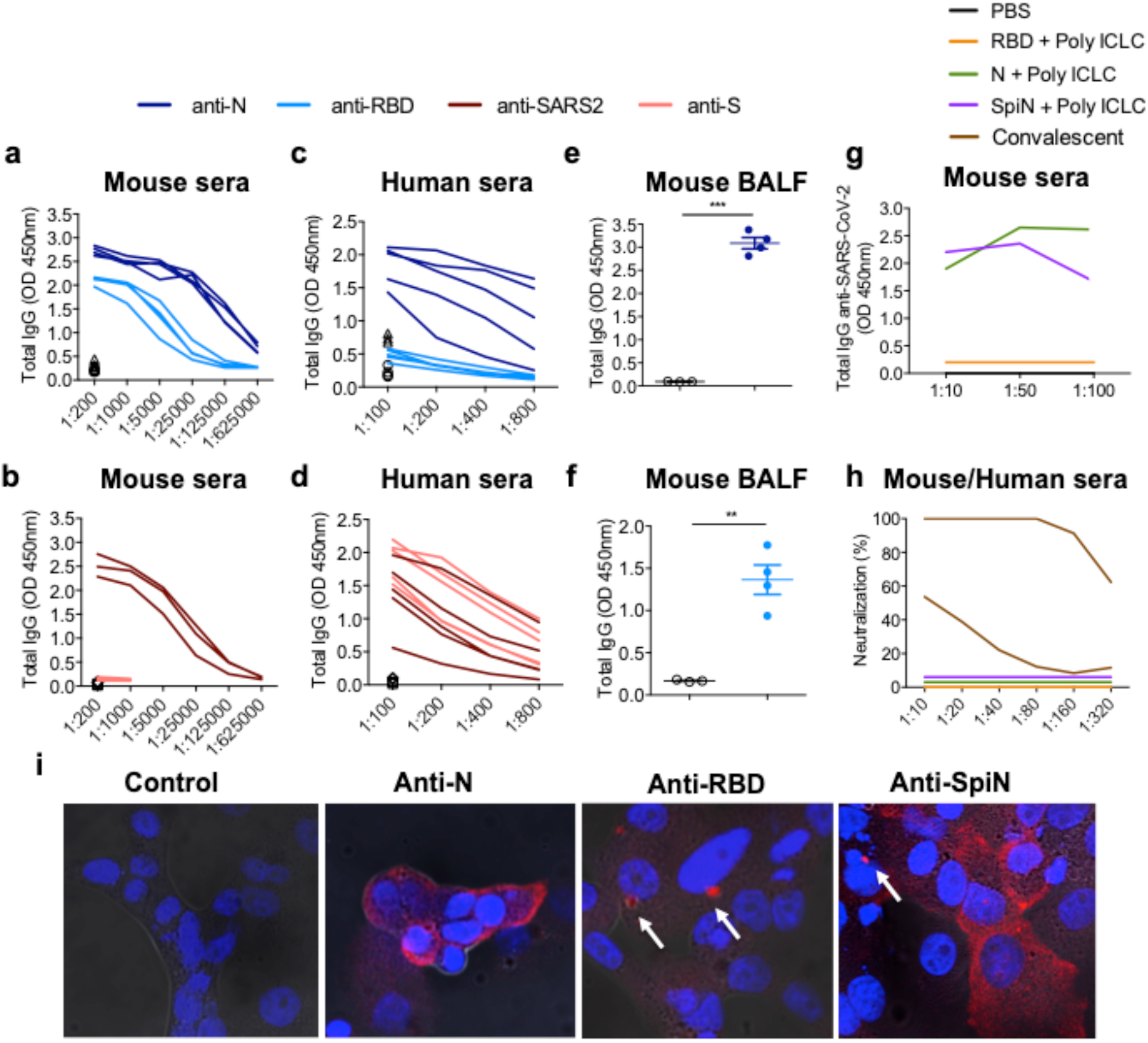
Levels of anti-SARS-CoV-2 total IgG and neutralizing antibodies in samples from mice immunized with SpiN. **a, b**, Total IgG responses of mice immunized with SpiN to N or RBD and inactivated SARS-CoV-2 and S (n=4 mice/group) **(b). c, d**, Total IgG response of convalescents to N or RBD and inactivated SARS-CoV-2 and S (n=5 mice/group) (d). **e, f**, The levels of anti-N **(e)** and anti-RBD **(f)** antibodies were measured in 1:1 diluted bronchoalveolar fluid (BALF) from mice that received Poly ICLC alone or SpiN associated with Poly ICLC (n=4 mice/group). **g, h**, The levels of anti-inactivated SARS-CoV-2 **(g)** and neutralizing antibodies **(h)** in pooled sera from mice immunized with either PBS (black line), N (green line), RBD (orange line) or SpiN (blue line) and from COVID-19 convalescent individuals (brown line). **i**, Immunofluorescence of SARS-CoV-2 infected cells stained with sera from mice that received adjuvant alone (left panel) or were immunized with either N (left middle panel), RBD (right middle panel) or SpiN (right panel) associated with Poly ICLC. Data are representative of two independent experiments. Statistical analysis of IgG measured in BALF was performed using two-tailed unpaired t-test, t=20.80 df=5 (anti-N), t=5.809 df=5 (anti-RBD). ** P < 0.01, *** P < 0.001.

Consistent with the high expression of the N protein ^27^ in infected cells, antibodies from mice immunized with either N or SpiN proteins, but not with the recombinant RBD, strongly recognized UV-inactivated SARS-CoV-2 in an ELISA (**Fig. 3g**). Relevant to this study, we found that immunization with neither N, RBD nor SpiN chimeric protein elicited nAbs to the wild type SARS-CoV-2, contrasting with the measurable levels in sera of COVID-19 convalescent patients (**Fig. 3h**). In agreement with the ELISA results (**Fig. 3g**), sera from mice immunized with the N protein strongly reacted with paraformaldehyde fixed SARS-CoV-2 infected cells showing a diffuse expression of N protein in the cytosol, as revealed by immunofluorescence (IFA) (**Fig. 3i**). In contrast, the sera from mice immunized with RBD anti-serum reacted with small vesicles in the cytosol, which might be consistent with Spike protein assembly in the endoplasmic reticulum–Golgi intermediate compartment (ERGIC) ^43^.The reactivity of sera from SpiN-immunized mice showed a mixed pattern consistent with a diffuse expression in the cytosol and a punctate staining, as observed with anti-N and anti-RBD antisera, respectively (**Fig. 3i)**.

As indicators of the immune response polarization, we measured the levels of IgG1 and IgG2c subclasses specific for N, RBD, S proteins and inactivated SARS-CoV-2 (**Fig. 4a-d and Extended Data Fig. 3b**,**c**). In mice immunized with SpiN associated to Poly ICLC, the levels of IgG2c antibodies to RBD are higher than IgG1, showing a trend to a type 1 immune response (**Fig. 4a**). In contrast, there is no difference in the levels of antigen specific IgG1 and IgG2c antibodies to N or inactivated SARS-CoV-2 (**Fig. 4b,c**). Similar levels of anti-RBD and anti-N of IgG1 and IgG2c in mice vaccinated with SpiN plus Addavax were observed (**Extended Data Fig. 4**). As shown for total IgG (**Fig. 3b**), the levels of antibodies to the S protein are very low (**Fig. 4d**). We also evaluated the recall T cell response in vaccinated mice, and despite the response to RBD was higher than the N, both proteins stimulated splenocytes to produce IFNγ (**Fig. 4e**). In contrast, they produced low levels of IL-10 when stimulated with SARS-CoV-2 proteins (**Fig. 4f**). Likewise, splenocytes from mice immunized with SpiN plus Addavax produced high levels of IFNγ and low levels of IL-10 (**Extended Data Fig. 3d**,**e**). The ELISPOT results were consistent with ELISA and showed a larger number of IFNγ-producing cells in response to RBD (**Fig. 4g**). In addition, flow cytometry shows that upon *in vitro* stimulation with proteins RBD and N, both CD4^+^ T and CD8^+^ T cells were highly activated as indicated by the expression of CD44 activation marker (**Fig. 4h,i**). When stimulated with soluble antigens, CD4^+^ T cells were shown to be the major source of IFNγ (**Fig. 4j,k**). Altogether, these results indicate that Poly ICLC induced a Type I-biased, whereas the use of Addavax as adjuvant yielded a mixed Type I/II immune response.

**Figure 4:**
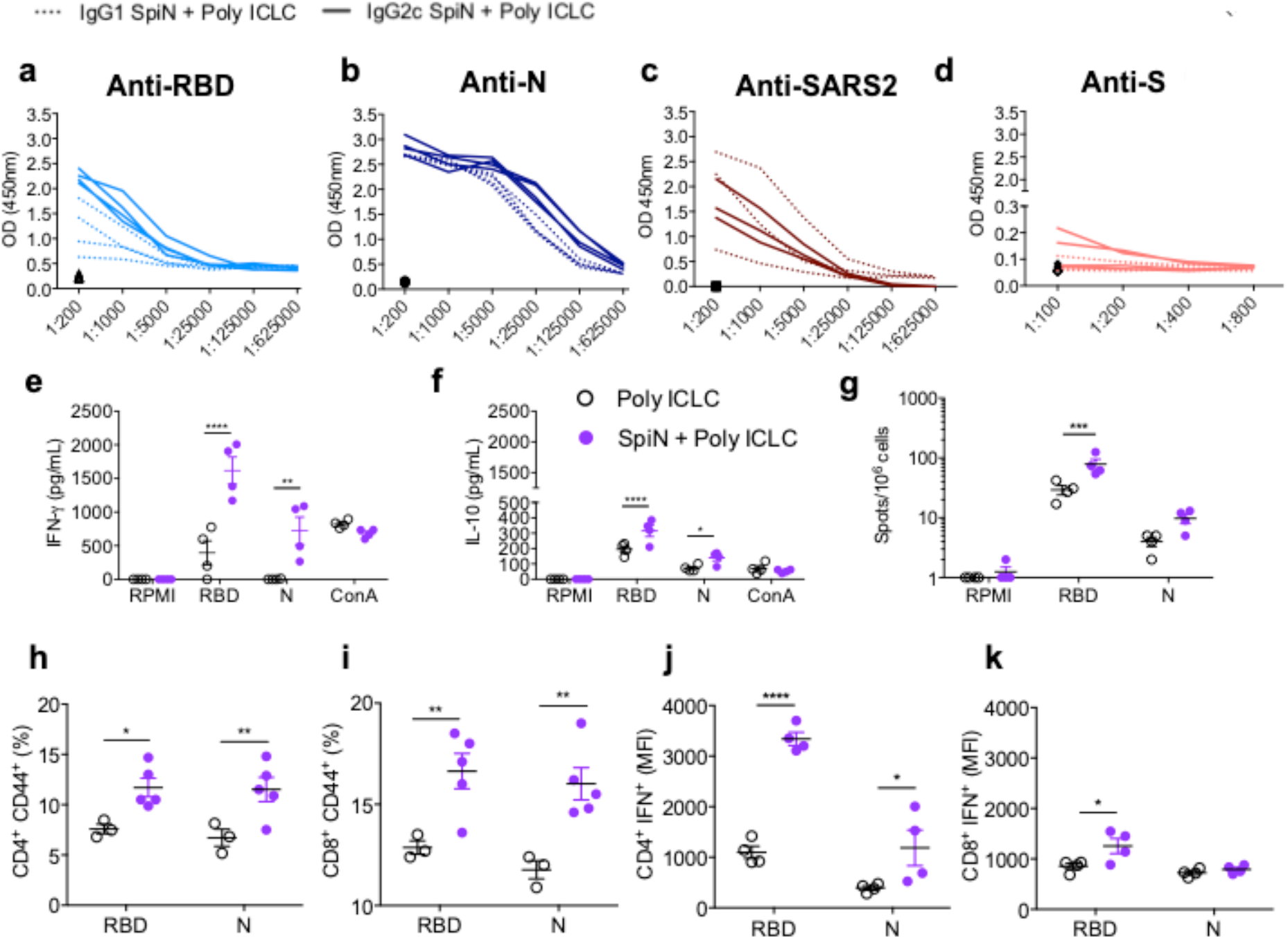
Levels of anti-SARS-CoV-2 IgG isotypes and T cell responses from mice immunized with SpiN associated with Poly ICLC. **a-d**, The levels of IgG1 and IgG2c specific for RBD **(a)**, N **(b)**, SARS-CoV-2 **(c)** and S **(d)** proteins in sera from mice that received adjuvant alone (black symbols in the bottom) or were immunized with SpiN protein associated with Poly ICLC. **e, f** The levels of IFNγ **(e)** or IL-10 **(f)** produced by splenocytes from vaccinated mice cultured in the absence (RPMI) or presence of RBD or N. Concanavalin A (ConA) was used as positive control. **g**, ELISPOT was used to quantify the frequency of IFNγ-producing T cells upon stimulation with either RBD or N protein. h-k, The frequency of activated CD4^+^ **(h)** and CD8^+^ **(i)** T lymphocytes was evaluated by measuring the expression of cell surface CD44, and intracellular expression of IFNγ **(j**,**k)** by flow cytometry. **a-k**, Data are representative of two independent experiments, n=4 mice/group. Statistical analysis of cytokine measurements and flow cytometry were performed using Two-way ANOVA followed by Sidak’s multiple comparisons test. **e**, df=24. **f**, df=36. **g**, df=18. **h**,**i** df=20. **j**,**k** df=12. * P < 0.05, ** P < 0.01, *** P < 0.001 and **** P < 0.0001.

Hamsters were used in our experiments as a model of moderate COVID-19 disease ^44^. The results presented in **Figure 5a** and **b** show that SpiN associated with either Poly ICLC or Addavax induced high levels of total IgG anti-N and anti-RBD in immunized hamsters. The levels of anti-RBD were relatively low in hamsters immunized with SpiN associated with Poly ICLC (**Fig. 5a,b**). As in SpiN immunized mice, the levels of nAbs in vaccinated hamsters were undetectable (**Fig. 5c**). Nevertheless, the viral load detect by RT-PCR were lower in vaccinated, as compared to unvaccinated hamsters challenged with SARS-CoV-2 (**Fig. 5d**). Histopathological analysis from challenged animals at 4 days post-infection (dpi) show that the lungs in the PBS group challenged with the Wuhan strain of SARS-CoV-2 revealed an accentuated diffuse alveolar wall thickening, with moderate multifocal collapse (asterisk), associated with accentuated diffuse congestion (black arrow) and mixed inflammatory infiltrate (mononuclear and polymorphonuclear cells). In the bronchial space, inflammatory cells are noted with a predominance of neutrophils associated with cellular debris (arrowhead) (**Fig. 5e, left panels**).

**Figure 5:**
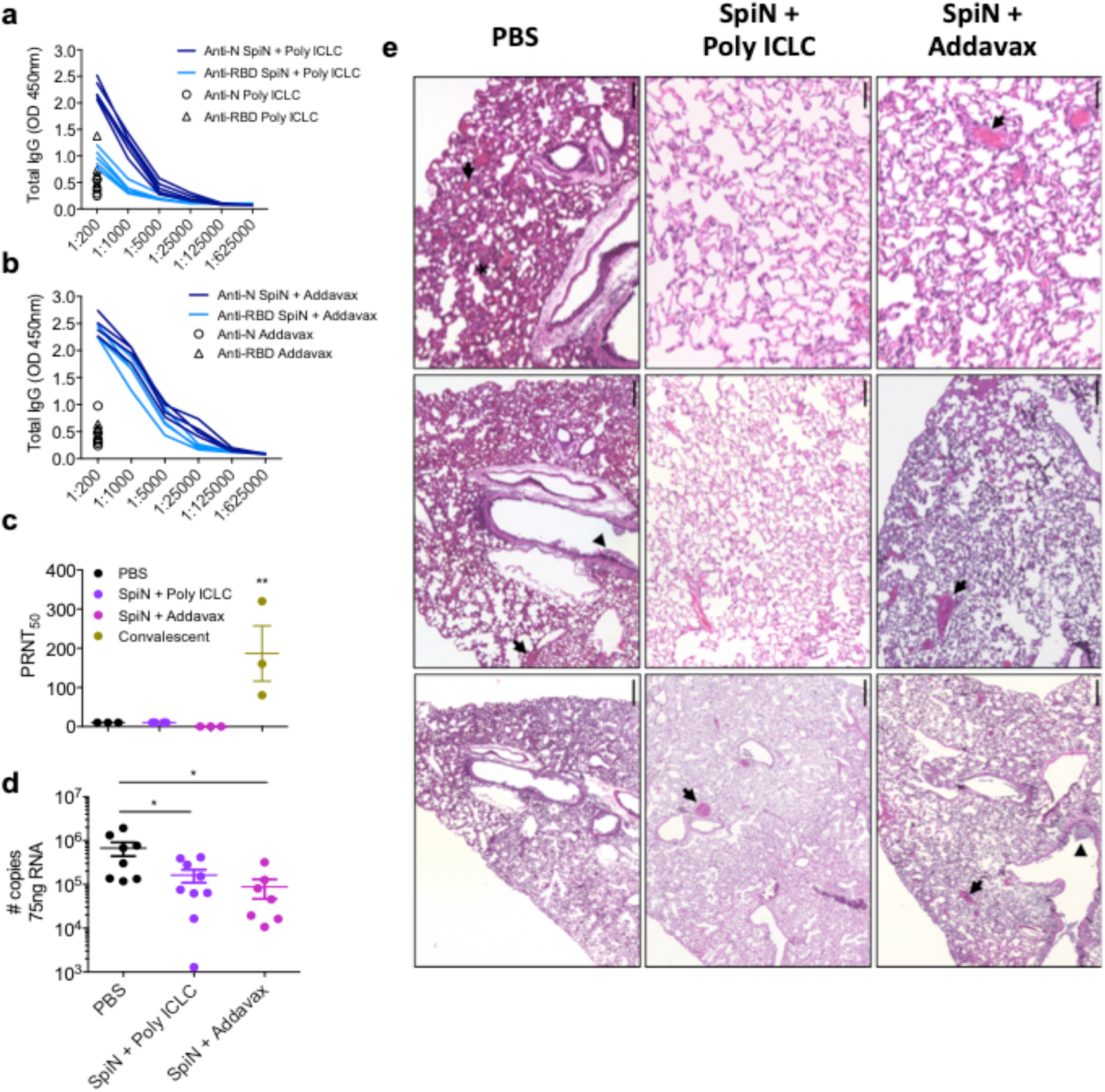
Protective immunity in hamsters immunized with SpiN and challenged with SARS-CoV-2. **a**,**b** The levels of total IgG anti-N and anti-RBD in sera of hamsters vaccinated with SpiN associated with Poly ICLC **(a)** and Addavax **(b). c**, Levels of neutralizing antibodies in the sera from hamsters vaccinated with SpiN + Poly ICLC or Addavax, in comparison with convalescent patients. **d**, Viral load measured by RT-PCR in the lungs of control and immunized hamsters with SpiN plus Poly ICLC or Addavax at 4 days post infection (DPI) with SARS-CoV-2. **e**, Histopathological sections of lungs from control and vaccinated hamsters at 4 DPI. The tissue was analyzed for congestion, inflammatory infiltrates, hemorrhagic foci, intra-alveolar exudate and alveolar collapse at 2,5x, 5x and 20x magnification. **a-c, e**, Data are representative of two independent experiments, n=5 hamsters/group. **d**, Pooled data from two independent experiments, n=8, 7 and 9 for PBS, SpiN + Poly ICLC and SpiN + Addavax, respectively. Statistical analysis of PRNT_50_ was performed using Kruskal-Wallis followed by Dunn’s multiple comparisons test. Data of viral load quantification was analyzed through One-way ANOVA followed by Bonferroni’s multiple comparisons test, df=21 * P < 0.05 and ** P < 0.01.

In comparison, hamsters immunized with SpiN adjuvanted with Poly ICLC had mild focal congestion (black arrow) with alveolar space preservation (**Fig. 5e, mid panels**). In the hamsters immunized with SpiN associated with Addavax moderate multifocal congestion is noted, associated with the presence of predominantly neutrophilic inflammatory infiltrate in the bronchi associated with cellular debris (arrowhead) (**Fig. 5e, right panels**).

Immunization with either SpiN associated to Poly ICLC or Addavax protected the K18-ACE-2 mice, a model of severe COVID-19, from weight loss (**Fig. 6a,e**) and other clinical signs of disease, such as affected motility, ruffle fur and hunching. Importantly, 100% of immunized mice survived until 11dpi, whereas all mice that received Poly ICLC or Addavax alone succumbed to infection (**Fig. 6b,f**). In addition, the viral RNA load both in the lungs and brain (**Fig. 6c,d**) were lower when comparing vaccinated (SpiN+Poly ICLC) versus non-vaccinated (Poly ICLC) controls, as indicated by RT-PCR. Similar results were obtained with mice immunized with SpiN adjuvanted with Addavax versus Addavax alone (**Fig. 6g-h**). Histopathological analysis demonstrate that the lungs from Poly ICLC group, at 5 dpi, showed a diffuse interstitial pneumonia characterized by a mixed inflammatory infiltration (mononuclear and polimorphonuclear cells), accompanied by intense congestion (black arrows), intra-alveolar exudate (white arrows with black outline), hemorrhagic foci (white star) and a few areas of alveolar collapse (asterisks) (**Fig 6i, top panels**). In comparison, the immunized group (SpiN plus Poly ICLC) showed preservation of the pulmonary architecture, with the presence of mononuclear peribronchovascular inflammatory infiltrate (red arrows) (**Fig. 6i, bottom panels**). Similar results were obtained with mice immunized with SpiN adjuvanted with Addavax. Of note, the control group (Addavax only) presented a diffuse and accentuated alveolar wall thickening (red arrowhead), with congestion (black arrows) associated with predominantly mononuclear inflammatory infiltrate with the presence of neutrophils (**Extended Data Fig. 3f, top panels**). In contrast, the immunized mice (SpiN plus Addavax) showed preserved lung architecture (**Extended Data Fig. 3f, bottom panels**). Consistent with the intense inflammatory response, we found high levels of mRNA expression of chemokines (CXCL2, CXCL5, CXCL9 and CXCL10) as well as IL-6 and TNFα in the lungs of infected mice. In contrast, the level of type 2 cytokines (IL-4 and IL-5) was higher in the vaccinated mice (**Figs. 6j,k**). These results indicate that cytokine mucosal environment was preserved in protected mice, whereas a dramatic switch to an inflammatory reaction was observed in animals that developed severe disease.

**Figure 6:**
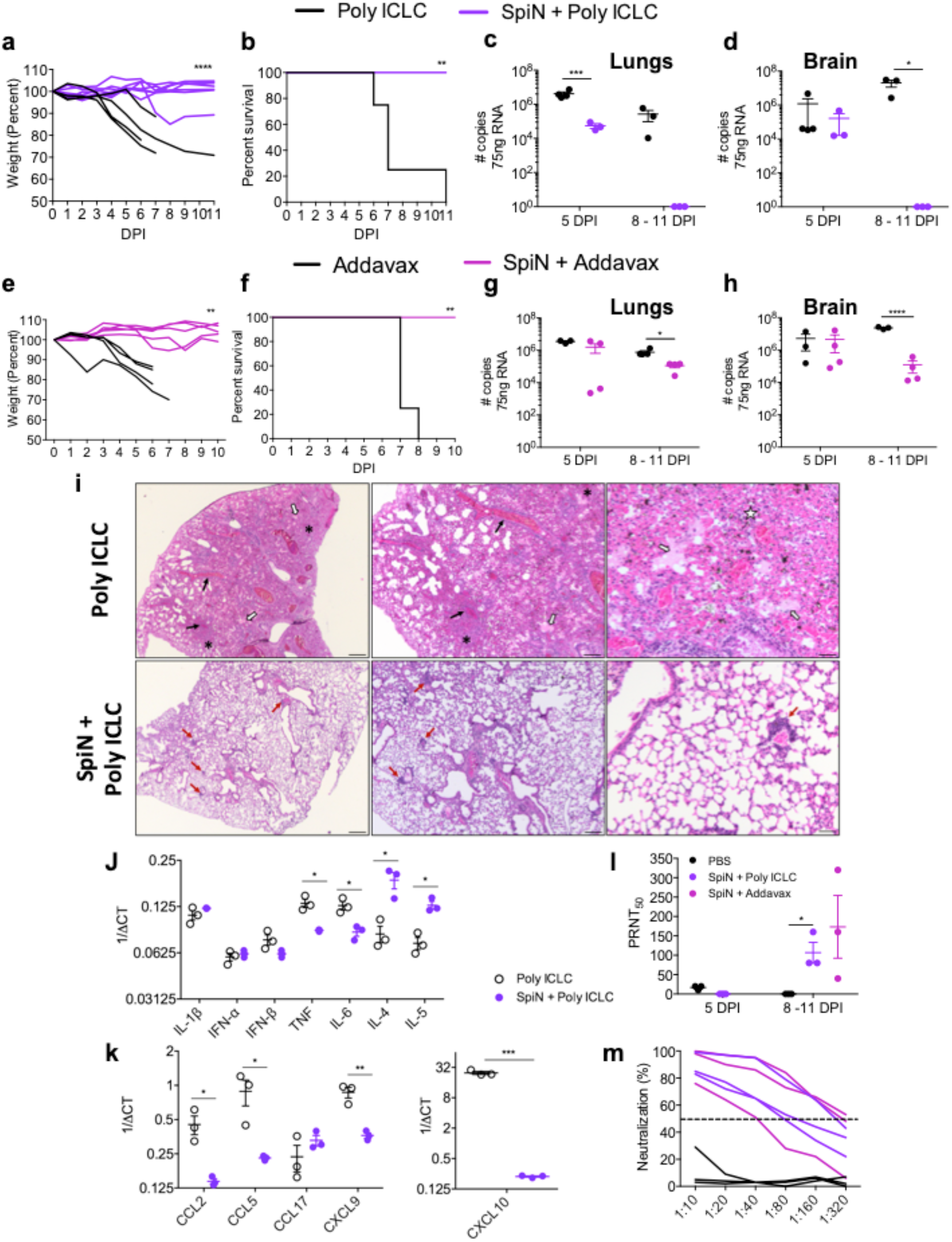
Protective immunity in mice immunized with the SpiN chimeric protein and challenged SARS-CoV-2. Mice were challenged with the Wuhan strain of SARS-CoV-2. Body weight **(a**,**e)** survival **(b**,**f)** and viral load in the lungs **(c**,**g)** and brains **(d**,**h)** from K18-hACE2 Tg mice immunized with SpiN associated with Poly ICLC or Addavax, and respective controls that received Poly ICLC or Addavax alone, as indicated. Viral load was measured by RT-PCR. **i**, Histopathological sections of lungs control and vaccinated hamsters at 5 and 7 days post-challenge with SARS-CoV-2. The tissue sections were stained with either hematoxylin and analyzed for congestion, inflammatory infiltrates, hemorrhagic foci, intra-alveolar exudate and alveolar collapse at 2,5x, 5x and 20x magnification. **j**,**k**, RNA was extracted from the lungs of control (Poly ICLC) and vaccinated (SpiN+Poly ICLC) mice at 5 dpi, reversely transcribed and the expression of cytokines **(j)** and chemokines **(k)** mRNAs was quantified by qRT-PCR. **l**,**m** nAbs on 5 or 8-11 DPI are shown as PRNT_50_ **(l)** and as percent of neutralization on 8-11 DPI **(m). a**,**b**,**e**,**f** Pooled data from two independent experiments. **a**,**b** n=4 for Poly ICLC and n=7 for SpiN + Poly ICLC. **e**,**f**, n=4 for Addavax, n=5 for SpiN + Addavax. **c**,**d**,**g-m** Data are representative of two independent experiments. **c**,**d**, n=3 mice/group. **g**,**h**, n=3 for Addavax and n=4 for SpiN + Addavax. **j-m**, n=3 mice/group. Statistical analysis of weight measurements was performed using Two-way ANOVA, df=89 **(a)**, df=49 **(e)**. Survival analysis was performed with Log-rank test. qRT-PCR data was analyzed with unpaired two-tailed t tests. Data of viral quantification was analyzed using Two-way ANOVA followed by Sidak’s multiple comparisons test, df=9 **(c**,**d)**, df=11 **(g)**, df=10 **(h)**. * P < 0.05, ** P < 0.01, *** P < 0.001 and **** P < 0.0001.

A relevant finding of this study is the demonstration that mice immunized with SpiN are highly resistant to SARS-CoV-2, even in absence of circulating neutralizing antibodies. An important question is regarding the levels of neutralizing antibodies in immunized rodents soon after the challenge with SARS-CoV-2. We found that at 4 dpi in hamsters (**Fig. 5c**) and 5 dpi in mice (**Fig. 6j**) the levels of neutralizing antibodies remained low, whereas the viral load was decreased in the lungs of immunized hamsters (**Fig. 5d**) and mice (**Fig. 6c,g**). This results further indicate a nAb-independent immunity in mice immunized with SpiN. At later time-points (8-11 days post-infection), we observed high levels of nAbs in sera of mice immunized with SpiN (**Fig. 6j,k**) and a drop in the viral load both in the lungs **(Fig. 6c,g)** and brain **(Fig. 6d,h**). These findings suggest that the priming of RBD-specific T cells during immunization may contribute to the secretion of nAbs triggered by the SARS-CoV-2 infection. We hypothesize that T cells act holding infection in the first days post-challenge and promote the production of nAbs at later stages of infection.

Since immunization did not induce any nAbs, we speculate that in this model, immunity at earlier stages of infection is solely mediated by T cells. However, vaccinated mice produced extremely high levels of anti-N antibodies and their role in mediating resistance to SARS-CoV-2 cannot be ruled out. Nevertheless, it was clear that the anti-N antibodies were uncapable of blocking *in vitro* invasion of host cell by SARS-CoV-2. While the N protein is associated with the viral genome, our immunofluorescence assay shows an overlap staining of the N protein with the surface membrane of the host cells. Thus, the anti-N antibodies might conceivably act by mediating antibody dependent cell cytotoxicity (ADCC) or by promoting internalization of opsonized host cells by macrophages. It is worth mentioning that infective SARS-CoV-2 does not grow well in macrophages, and we have no evidence that anti-N antibodies bind to viral particles and promote antibody-dependent enhancement (ADE) *in vitro*, or enhanced *in vivo* SARS-CoV-2 replication in tissues from vaccinated mice.

In conclusion, immunization with the N and RBD fusion protein adjuvanted with either Poly-ICLC or Addavax was highly efficient in protecting against SARS-CoV-2 challenge. Altogether our results support the hypothesis that this is primarily due to T cell-mediated immunity. Since T cell responses do not rely on a single epitope, it is unlikely that non-silent point mutations will severely undermine immune-mediated resistance induced by vaccination with the SpiN chimeric protein. Hence, while not denying the importance of nAbs, the N protein and more broadly the use of multiple T cell epitopes, should be considered to improve anti-COVID-19 vaccines and for the development of vaccines that overcome SARS-CoV-2 genetic plasticity.

## Methods

### Blood donors and ethics statement

Blood samples were collected from vaccinated individuals (n=33), convalescent patients (n=13), and healthy controls (n=9). All individuals were between 18 and 70 years old (36±11, female:male ratio=3.2) (**Extended Table I**). Vaccinated individuals received two doses of Coronavac (Sinovac, China) and were sampled 27-54 days after the second dose. Convalescent individuals reported having mild COVID-19 between 24-196 days before sampling, confirmed by PCR. All individuals were briefly interviewed before sampling and consent forms were signed. This study was performed under protocols reviewed and approved by the Ethical Committees on Human Experimentation from Fundação Hospitalar do estado de Minas Gerais (FHEMIG) - CAAE: 43335821.4.0000.5119. All patients were adults and were enrolled in the study after providing written informed consent.

### Mice, hamsters, virus and ethics statement

Female C57BL/6 mice, 6-10 weeks old, were purchased from the Center for Laboratory Animal Facilities of the Federal University of Minas Gerais (CEBIO-UFMG). Human Angiotensin Converting Enzyme transgenic mice (K18-hACE2) mice in the C57BL/6 background, originally from Jackson Laboratories, were bred at Fiocruz-Minas or at Fiocruz-São Paulo animal facilities and used as a model of severe COVID-19. Female Golden Syrian Hamsters, 6-10 weeks old were from Fiocruz-Minas Animal House and used as a model of mild COVID-19. The severe acute respiratory syndrome coronavirus 2 (SARS-CoV-2) viral strain used in this study was from the lineage B (isolate BRA/SP02/2020). Viral stocks were propagated and titrated in Vero E6 cells (ATCC CRL-1586) by plaque assay according to previous standardized conditions ^45^. Viral aliquots were kept in -80ºC until further use. The experiments were carried out following the recommendations of the Guide for the Care and Use of Laboratory Animals of the Brazilian National Council of Animal Experimentation (CONCEA). The protocols for animal experiments were approved by Fundação Oswaldo Cruz and Universidade Federal de São Paulo Ethics. Commission on Animal Use (CEUA) LW 25/20 and 105/2020, respectively.

### Epitope prediction and sites of amino acid changes in the RBD region of the Spike (S) and Nucleocapsid (N) proteins from SARS-CoV-2

Epitope prediction was performed through The Immune Epitope Database and Analysis Resource (IEDB) platform for class I analysis and NetMHCII for class II ^30,31^. Potential HLA-ABC, HLA-DR and mice MHC-I/II binding epitopes were sought in the sequences of Spike (accession BCN86351) and Nucleocapsid (accession QRV71356) proteins. Isolates from the five most relevant groups of SARS-CoV-2 variants GISAID (UK - VUI202012 / 01 GRY-B.1.1.7; South Africa - GH/501Y.v2-B.1.351; Brazil - GR/501Y.V3-P.1; USA-Ca - GH/452R. V1-B.1.429 + B.1.427; UK/Nigeria - G/484K.V3-B.1.525) were analyzed. The sequences were translated *in silico* using the standard genetic code and multiple aligned together with N or S proteins of Lineage B (Wuhan – EPI_ISL_402123) by the Kalign tool ^46^ with parameters 8.52 “gap extension penalty”, 54.90 “gap open penalty” and 4.42 “gap terminal penalty”. To build the Needle plot, a “homemade” script was implemented to count the number of divergences per alignment position considering each of the translated sequences in relation to the sequences of S or N proteins from lineage B. The number of amino acid divergences in each position was normalized by the number of N or S proteins initially selected for the analysis. The R package “mutsneedle” was used to construct the figure. The dendrograms was constructed using the iqtree tool ^47^, employing the “Maximum likelihood” statistical approach and the JTTDCMut+I substitution model to obtain the correlations from the N or S amino acids sequences aligned. Lineage B was defined as “outgroup” and the R ggtree package ^48^ was used to the tree visualization.

### Plasmid constructions and recombinant antigens production

Plasmids containing sequences encoding the full length N, the RBD of Spike and the quimeric SpiN protein with codons optimized for expression in *E. coli* were purchased from Genscript. Competent *E. coli* Star™ (DE3) were transformed with the pET24 vector with N or the RBD sequences and *E. coli* pRARE with the pET24_with SpiN. Transformed bacteria were grown in LB medium with kanamycin (50 μg/ml) at 37°C until OD600 0.6 was reached. At this point, protein expression was induced by adding IPTG to the culture at a final concentration of 0.5 mM. The induction of expression of the three proteins was done at 37°C for 3h for N and RBB and for 18h for SpiN. The N and RBD proteins contained a histidine tag and were purified through affinity chromatography step with the Histrap HP (GE HealthCare) column following the manufacturer’s instructions. After bacterial lysis, N protein was purified from the soluble fraction and RBD protein from the insoluble fraction by adding 8M urea in the buffers for solubilization. The SpiN protein, expressed without histidine tag, was purified from the soluble and insoluble fractions of the bacteria cell lisate after addition of 8M urea and through two steps of chromatography. A cation exchange chromatography with the Hitrap SP HP column (GE HealthCare) was followed by molecular exclusion with the column HiPrep 26/60 Sephacryl S-100 HR (GE HealthCare), following the manufacturer’s instructions. The S protein expressed in mammalian cells was kindly provided by Dr. Leda Castilho from Universidade Federal do Rio de Janeiro.

### PBMC cultures and IFNγ measurements

Peripheral blood mononuclear cells (PBMCs) were isolated from heparinized blood by ficoll gradient. Briefly, blood layered on Ficoll-Paque plus (GE-Healthcare) were centrifuged 410 x g, 40 minutes, RT. One-million cells per well were distributed in 96-well flat-bottom plates and incubated in complete media (RPMI 1640, 10% FBS, 100 mg/ml streptomycin, 100 U/ml penicillin) with 5 μg/mL from either N, RBD and S recombinant antigen or anti-CD3 (1μg/mL) and anti-CD28 (0.5μg/mL) as positive controls. Unstimulated cells were used to assess the background production of cytokines. Culture supernatants were harvested after 72h and frozen at -80°C until analysis. Levels of IFNγ were measured by ELISA following the manufacturer’s protocol (BD, OptEIA™ Human IFNγ).

### Detection of human, mouse and hamster antigen-specific antibodies

Plates were coated overnight with 0.4 μg/well of either N, RBD, S recombinant proteins, or alternatively 10^4^ PFU/well of UV-inactivated SARS-CoV-2, and blocked for 2 hours with PBS containing 2% bovine serum albumin (PBS-2% BSA) at 37°C. Serum were serially diluted and the bronchoalveolar lavage (BALF) samples tested at 1:1 dilution in PBS-2% BSA and incubated for 1 hour at 37°C, and then incubated with anti-human IgG-HRP antibody (Fapon), anti-hamster IgG-HRP or anti-mouse total IgG, IgG1, IgG2c conjugated with streptavidin-HRP (Southern Biotech). After 5 washes, plates were revealed with 1-Step ultra TMB substrate solution (Biolegend) for 15 minutes in the dark, and reaction was stopped by adding 2N H_2_SO_4_ (Sigma). Plates were read in 450nm and results were expressed as raw optical density (OD). The antibody titer was determined by the sera dilution that yielded 50% of the maximum antibody reactivity to the antigen in the ELISA.

### Reduction Neutralization Assay

One day prior infection, 10^5^ Vero E6 cells were seeded in Dulbecco’s modified eagle media (DMEM) (Vitrocell, Brazil) with 10% Fetal Bovine Serum (FBS) to each well of a 48 wells plate. On the next day, sera samples from mice or human were heat inactivated by incubation at 56ºC during 1h on a warm bath. Samples were two-fold serially diluted (1:10 to 1:320 (v/v)) in DMEM and mixed with 100 PFUs of SARS-CoV-2 viral stock. Media only was used instead of sera sample for positive control. The mixture was incubated during 1h ta 37ºC to allow antibody binding to viral particle. Next, Vero E6 cell culture supernatant was removed and the cells were inoculated with 50μl/well of sera-virus mixture, incubated for 1h at room temperature under gently rocking to allow viral-biding to cells. Then, 1ml of pre-warmed DMEM with 2% FBS and 2% of carboxymethylcellulose (CMC) was gently added to each well and the plates were incubated at 37ºC and 5% CO_2_ atmosphere during 4 days to allow viral plaque formation. Then, the cells were fixed with 4% formaldehyde solution diluted in PBS during 2 hours and stained with 1% Naphtol blue black (Sigma, USA) solution for 1 hour for plaque visualization. Neutralization activity were determined by plaque numbers reduction compared to the positive control.

### Viral Quantitative Reverse Transcription Polymerase Chain Reaction

Total RNA was extracted from homogenized mice tissues using the RNA Mini Kit (Qiagen, USA), according to protocols provided by the manufacturers. qRT-PCR was performed in 12 μL reactions using GoTaq Probe 1-step RT-qPCR System (Promega, US) according to the manufacturer’s instructions, using 75 ng of total RNA per reaction. Primers and fluorescente probes were designed based on previous described diagnostic qRT-PCR protocol specific for SARS-CoV-2, which amplify a 100bp amplicon from the E gene of SARS-CoV-2 ^49^. Cycling conditions were 45ºC for 15 min and 95ºC for 3 min followed by 45 cycles at 95ºC for 15s and 58ºC for 60s, using Quantstudio 5 Real Time PCR system (Applied Biossystems, USA). For viral load quantification, a standard curve based on plasmid containing the E gene sequence (SARS-CoV-2 Wuhan-Hu isolate sequence) was constructed. Serial 10-fold dilutions of plasmid DNA that correspond to viral copies ranging from 2 to 2 × 10^5^ were used as templates to prepare the standard curves. Real-time PCR assays were carried out in triplicate and the resulting Ct values by plotted against copy number of viral genome.

### Immunization, challenge and histopathology

Hamsters and C57BL/6 or hACE2 mice received two administrations at 21 days apart, containing 10 μg of RBD, N or SpiN adjuvanted with 50 μg of Hiltonol® (Poly ICLC, supplied by Oncovir, Washington, D.C.) ^39,40^, or with AddaVax™ (oil-in-water emulsion similar to MF59) in a volume of 1:1 AddaVax™:Antigen ^50^. The solution was inoculated intramuscularly in a final volume of 50 μL into each tibial muscle. Thirty days post immunization, animals were challenged intranasally with 5 × 10^4^ PFU of SARS-CoV-2 (for hACE2 mice) and 10^5^ PFU for hamsters. The body weight, clinical signs and survival were evaluated for 11 days post infection. For the histopathology analyses, harvested tissues were fixed in phosphate-buffered 10% formalin for seven days, embedded in paraffin, processed using a Tissue processor PT05 TS (LUPETEC, UK) tissue processor and embedded in histological paraffin (Histosec®, Sigma-Aldrich). The 4 μm thick sections were stained with hematoxylin and eosin.

### Measurements of mouse cytokines

Mouse splenocytes were isolated by macerating the spleen through a 100 μm pore cell strainer (Cell Strainer, BD Falcon) followed by treatment with ACK buffer for erythrocytes lysis. The number of cells was adjusted to 10^6^ cells per well and then stimulated with 10 μg/mL of RBD or N. Concanavalin A (Sigma, 5 μg/mL) was used as positive control. The supernatants were collected 72 hours post-stimulation and the levels of IFNγ and IL-10 determined by and ELISA (R&D Systems). IFNγ production was also assessed by ELISPOT, in which plates (MAHAS4510 – Millipore) were coated with capture antibody anti-IFN-γ (R4-6A2 - BD) and incubated overnight at 4°C. The plates were blocked and 10^6^ cells were added per well, together with 10 μg/mL of RBD or N stimuli plus rIL-2 (100 U/mL) and anti-CD28 (1 μg/mL). After 20 hours of culture, the plates were washed and incubated with biotinylated antibody anti-IFN-γ (XMG1.2 – BD). The reaction occurred after the addition of streptavidin-HRP for 1 hour followed by incubation with 3,3’-Diaminobenzidine (DAB).

### Cytokines and chemokines measurements by qRT-PCR

The RNA samples isolated from the lungs of immunized and challenged mice at 5DPI were treated with DNase (Promega), and then converted into cDNA using High-Capacity cDNA Reverse Transcription Kit (Applied Biosystems), according to manufacturer’s instructions. qPCRs reactions were performed with Sybr Green PCR Master Mix (Applied Biosystems) in an ABI7500 Real Time PCR System (Applied Biosystems), under standard conditions. Primer sequences are presented in **Extended Table 2**. qRT-PCR data were presented as 1/ΔCT.

### Immunofluorescence assays

A 16-well chamber slide (ThermoFisher) was coated with 10^4^ Vero E6 cells/well and incubated overnight with SARS-CoV-2 in a multiplicity of infection (M.O.I) of 10. Then, the cells were treated with Brefeldin A for 4 hours, fixed with paraformaldehyde 4% and permeabilized with PBS-P (PBS 0,5% BSA + 0,5% saponin). Later, the wells were blocked with BSA 1% and incubated with sera from mice immunized with RBD, N or SpiN. The secondary antibody Alexa Fluor 594 anti-mouse IgG (ThermoFisher) was added and the nucleus was stained with DAPI (ThermoFisher). The slides were analyzed in the confocal microscope LSM 780 Carl Zeiss AxioObserver, objective 63x oil NA 1.4. Images were processed with the software ImageJ version 2.1 for Mac.

### Flow cytometry

For imunophenotyping, a total of 2 × 10^6^ of splenocytes derived from immunized mice were incubated for 18h at 37°C and 5% CO_2_ with RPMI 1640 medium alone or containing 10 μg/mL of RBD or N proteins. During the last 6 hours of culture, GolgiStop and GolgiPlug Protein Transport Inhibitors (BD Biosciences) were added to the cell cultures. The splenocytes were then washed with PBS, stained with Live/Dead reagent (Invitrogen) and incubated with FcBlock (BD Biosciences) ^51^. The following mAbs were used to label cell surface markers: anti-CD3 PerCP-Cy5.5 (BD, clone 145-2C11), anti-CD4 Alexa Fluor 700 (Invitrogen, clone KG1.5), anti-CD8 APC-Cy7 (Biolegend, clone 53-6.7) and anti-CD44 APC (Invitrogen, clone IM7). For intracellular staining, cells were washed, fixed and permeabilized according to the manufacturer’s instructions (Cytofix/Cytoperm, BD Biosciences) and stained with anti-IFNγ APC (BD, clone XMG1.2). Flow cytometry was carried out using a BD LSRFortessa and ∼100,000 live CD3^+^ cells were acquired. Data were analyzed using FlowJo v10.5.3 software.

### Statistical analysis

Statistical analysis was conducted using GraphPad Prism 6.0 for Mac (GraphPad Inc, USA). First, outliers were detected with Grubbs’s test and then D’Agostino-Pearson was run to verify data normality. The tests used on each data analysis are explained on figure legends. In general, comparison between the groups were performed through unpaired t-test or Mann–Whitney U test, according to data distribution. Weight measurement data were analyzed by Two-Way ANOVA, followed by Sidak’s multiple comparisons test. For survival analysis, the log-rank test was used. Statistical differences were considered significant when p values ≤ 0.05.

### Reporting summary

Further information on research design is available in the Nature Research Reporting Summary linked to this paper.

## Data availability

The authors declare that all data supporting the findings of this study are available within the paper and its supplementary information files. If any more information is needed, data are available from the corresponding author upon reasonable request.

## Acknowledgements

This work was funded by Rede Virus from the Ministry of Science, Technology and Innovation (Finep 01.20.0010, 01.20.0005.00, 0069/15); CNPq 403514/2020-7 and 403701/2020-1), National Institute of Science and Technology of Vaccines (Fapemig APQ-03608-17; CNPq 465293/2014-0), FAPESP 2020/05527-0), Ministry of Education (CAPES), Fundação Oswaldo Cruz (INOVA, 2020), Prefeitura de Belo Horizonte, and Parliamentary Amendment of State and Federal Congress Representatives from Minas Gerais. We thank Dr. Luis Adam and Dr. Flávia Bagno for providing the sera from convalescent COVID-19 patients and our project analyst Ms. Cristiane Gomes.

## Author Contributions

J.C. performed epitope predictions. R.M. was responsible for the construction of the needle plot and dendograms. J.C., G.B. and D.D. contributed to protein’s conception and plasmids constructions. N.S. and B.C. performed protein expression and purification, SDS-PAGE and Western Blots. G.G.A., L.I.O., T.G.M. and M.A. performed IgG and IFN-γ ELISAS of human samples. J.C., N.S.H-S and P.A. were responsible for mice and hamsters immunizations and ELISAS. J.C. and N.S.H-S performed ELISPOT and flow cytometry. J.C., M.J.F. and P.A. performed the challenge experiments with SARS-CoV-2. M.J.F. performed the neutralization assays. J.C. and M.J.F. performed immunofluorescence assays. J.C., M.J.F. and L.F. performed viral RNA extraction and quantification by RT-PCR. J.C. performed cytokine and chemokine mRNA measurements by RT-PCR. B.R. and S.G.R. performed histopathological analysis. E.D. provided SARS-CoV-2 viruses. A.S. and O.C. provided reagents. J.C., M.J.F., R.T.G, S.R.T., A.P.F., F.F., J.S.S., A.M. J.K. and E.D. contributed to study’s conception and design. J.C. and R.T.G. wrote the manuscript. All authors discussed the results and the manuscript.

## Additional Information

**Supplementary Information** is available for this paper.

**Correspondence and requests for materials** should be addressed to Ricardo T. Gazzinelli.

**Reprints and permissions information** is available at http://www.nature.com/reprints.

