## Supplementary Data for "Neutralizing antibody-independent immunity to SARS-CoV-2 in hamsters and hACE-2 transgenic mice immunized with a RBD/Nucleocapsid fusion protein"

### Extended Data

**Extended Table I -** Healthy donors, COVID-19 convalescent and CoronaVac vaccinated individuals enrolled in our study.

| Cod | Group | Sex | Age | Days between vaccination and sampling | Sampling date | Vaccination day (2nd dose) | Date of COVID PCR+ | Vaccine |
|---|---|---|---|---|---|---|---|---|
| **804-R** | Conv | Female | 22 | NA | 03/25/21 | NA | 09/05/20 | NA |
| **805-R** | Conv | Female | 29 | NA | 03/25/21 | NA | 03/01/21 | NA |
| **806-R** | Conv | Female | 20 | NA | 03/25/21 | NA | 01/09/21 | NA |
| **807-R** | Conv | Female | 33 | NA | 03/30/21 | NA | 01/15/21 | NA |
| **808-R** | Conv | Male | 23 | NA | 04/01/21 | NA | 03/10/21 | NA |
| **809-R** | Conv | Female | 44 | NA | 04/01/21 | NA | 12/22/20 | NA |
| **810-R** | Conv | Female | 39 | NA | 04/01/21 | NA | 11/21/20 | NA |
| **811-R** | Conv | Female | 32 | NA | 04/06/21 | NA | 03/20/21 | NA |
| **812-R** | Conv | Female | 38 | NA | 04/06/21 | NA | 12/05/20 | NA |
| **813-R** | Conv | Male | 26 | NA | 04/06/21 | NA | 09/22/20 | NA |
| **814-R** | Conv | Female | 59 | NA | 04/27/21 | NA | 03/22/21 | NA |
| **817-R** | Conv | Female | 25 | NA | 05/04/21 | NA | 04/08/21 | NA |
| **820-R** | Conv | Female | 35 | NA | 05/04/21 | NA | 04/02/21 | NA |
| **038-0** | HD | Female | 40 | NA | 04/06/21 | NA | 06/15/20 | NA |
| **063-0** | HD | Female | 60 | NA | 05/07/21 | NA | NA | NA |
| **104-0** | HD | Female | 18 | NA | 06/26/21 | NA | NA | NA |
| **105-0** | HD | Male | 25 | NA | 06/24/21 | NA | NA | NA |
| **904-H** | HD | Female | 20 | NA | 03/25/21 | NA | NA | NA |
| **906-H** | HD | Male | 28 | NA | 03/25/21 | NA | NA | NA |
| **908-H** | HD | Male | 48 | NA | 03/30/21 | NA | NA | NA |
| **909-H** | HD | Male | 38 | NA | 03/30/21 | NA | NA | NA |
| **913-H** | HD | Female | 38 | NA | 04/06/21 | NA | NA | NA |
| **001-A** | Vacc | Female | 25 | 33 | 03/16/21 | 02/11/21 | NA | Coronavac |
| **002-A** | Vacc | Female | 27 | 33 | 03/16/21 | 02/11/21 | NA | Coronavac |
| **003-A** | Vacc | Female | 26 | 30 | 03/16/21 | 02/14/21 | NA | Coronavac |
| **004-A** | Vacc | Male | 56 | 32 | 03/16/21 | 02/12/21 | NA | Coronavac |
| **005-A** | Vacc | Male | 31 | 27 | 03/16/21 | 02/17/21 | NA | Coronavac |
| **006-A** | Vacc | Female | 39 | 29 | 03/18/21 | 02/17/21 | NA | Coronavac |
| **007-A** | Vacc | Female | 28 | 29 | 03/18/21 | 02/17/21 | NA | Coronavac |
| **008-A** | Vacc | Female | 27 | 29 | 03/18/21 | 02/17/21 | NA | Coronavac |
| **009-A** | Vacc | Female | 30 | 28 | 03/18/21 | 02/18/21 | NA | Coronavac |
| **010-A** | Vacc | Female | 36 | 31 | 03/18/21 | 02/15/21 | 08/15/20 | Coronavac |
| **017-A** | Vacc | Female | 27 | 41 | 03/25/21 | 02/12/21 | NA | Coronavac |
| **018-A** | Vacc | Female | 26 | 45 | 03/25/21 | 02/08/21 | NA | Coronavac |
| **019-A** | Vacc | Female | 27 | 38 | 03/25/21 | 02/15/21 | NA | Coronavac |
| **020-A** | Vacc | Female | 37 | 38 | 03/25/21 | 02/15/21 | NA | Coronavac |
| **021-A** | Vacc | Female | 39 | 48 | 03/25/21 | 02/05/21 | NA | Coronavac |
| **022-A** | Vacc | Female | 70 | 34 | 03/25/21 | 02/19/21 | 03/28/20 | Coronavac |
| **023-A** | Vacc | Female | 31 | 36 | 03/30/21 | 02/22/21 | NA | Coronavac |
| **024-A** | Vacc | Female | 38 | 47 | 03/30/21 | 02/11/21 | NA | Coronavac |
| **025-A** | Vacc | Female | 39 | 46 | 03/30/21 | 02/12/21 | NA | Coronavac |
| **026-A** | Vacc | Female | 48 | 43 | 03/30/21 | 02/15/21 | NA | Coronavac |
| **027-A** | Vacc | Female | 50 | 48 | 03/30/21 | 02/10/21 | NA | Coronavac |
| **028-A** | Vacc | Female | 54 | 42 | 03/30/21 | 02/16/21 | NA | Coronavac |
| **029-A** | Vacc | Female | 42 | 42 | 03/30/21 | 02/16/21 | NA | Coronavac |
| **030-A** | Vacc | Female | 28 | 43 | 03/30/21 | 02/15/21 | NA | Coronavac |
| **031-A** | Vacc | Male | 60 | 44 | 04/01/21 | 02/16/21 | NA | Coronavac |
| **034-A** | Vacc | Male | 47 | 45 | 04/01/21 | 02/15/21 | NA | Coronavac |
| **035-A** | Vacc | Male | 39 | 45 | 04/01/21 | 02/15/21 | 11/15/20 | Coronavac |
| **036-A** | Vacc | Female | 44 | 43 | 04/01/21 | 02/17/21 | NA | Coronavac |
| **037-A** | Vacc | Female | 28 | 47 | 04/06/21 | 02/18/21 | 11/26/20 | Coronavac |
| **039-A** | Vacc | Female | 29 | 53 | 04/06/21 | 02/12/21 | 11/27/20 | Coronavac |
| **040-A** | Vacc | Female | 27 | 54 | 04/06/21 | 02/11/21 | NA | Coronavac |
| **041-A** | Vacc | Male | 60 | 54 | 04/06/21 | 02/11/21 | NA | Coronavac |
| **042-A** | Vacc | Male | 31 | 49 | 04/06/21 | 02/16/21 | NA | Coronavac |

Abbreviations: NA, not applicable; Conv, convalescent; HD, healthy donors; Vacc, vaccinated.

**Extended Table II –** Primer sequences used to quantify cytokine and chemokine mRNAs by RT-PCR.

| Gene | Primer Sequences | |
|---|---|---|
| IL-1β | F 5’ ACCTGTCCTGTGTAATGAAAGACG 3’ | R 5’ TGGGTATTGCTTGGGATCCA 3’ |
| IFN-α | F 5’ GTTCAAGTCTCTGTCCCCAAAA 3’ | R 5’ GTGGGAACTGCACCTGATGT 3’ |
| IFN-β | F 5’ CAGCTCCAAGAAAGGACGAAC 3’ | R 5’ GGCAGTGTAACTCTTCTGCAT 3’ |
| TNF | F 5’ CCCTCACACTCAGATCATCTTCT 3’ | R 5’ GCTACGACGTGGGCTACAG 3’ |
| IL-6 | F 5’ TGTTCTCTGGGAAATCGTGGA 3’ | R 5’ AAGTGCATCATCGTTGTTCATACA 3’ |
| IL-4 | F 5’ TCATCGGCATTTTGAACGAG 3’ | R 5’ CGTTTGGCACATCCATCTCC 3’ |
| IL-5 | F 5’ AAAGAGAAGTGTGGCGAGGAGA 3’ | R 5’ CACCAAGGAACTCTTGCAGGTAA 3’ |
| CCL2 | F 5’ TGGCTCAGCCAGATGCAGT 3’ | R 5’ TTGGGATCATCTTGCTGGTG 3’ |
| CCL5 | F 5’ CAAGTGCTCCAATCTTGCAGTC 3’ | R 5’ TTCTCTGGGTTGGCACACAC 3’ |
| CCL17 | F 5’ CAGGGATGCCATCGTGTTTC 3’ | R 5’ CACCAATCTGATGGCCTTCTT 3’ |
| CXCL9 | F 5’ AATGCACGATGCTCCTGCA 3’ | R 5’ GGTCTTTGAGGGATTTGTAGTG 3’ |
| CXCL10 | F 5’ GCCGTCATTTTCTGCCTCA 3’ | R 5’ CGTCCTTGCGAGAGGGATC 3' |

### Extended Data Fig. 1

**a**

| | Peptide | HLA-ABC |
|---|---|---|
| RBD | KLNDLCFTNV | 02:01 |
| | FELLHAPATV | 02:01; 40:01 |
| | KLPDDFTGCV | 02:01 |
| | LYNSASFSTF | 24:02 |
| | YNSASFSTFK | 11:01 |
| | NLDSKVGGNY | 01:01 |
| | RQIAPGQTGK | 03:01; 11:01 |
| | LPIGINITRF | 07:02 |
| | QPYRVVVLSF | 07:02 |
| | IVRFPNITNL | 08:01 |
| | FERDISTEIY | 44:02 |
| | TESNKKFLPF | 44:02 |
| Nucleoprotein | YLGTGPEAGL | 02:01 |
| | WLTYTGAIKL | 02:01 |
| | LLLLDRLNQL | 02:01; 08:01 |
| | ADLDDFSKQL | 02:01; 40:01; 44:02 |
| | VLQLPQGTTL | 02:01; 40:01; 44:02; |
| | FGMSRIGMEV | 02:01 |
| | QFAPSASAFF | 24:02 |
| | AQFAPSASAF | 24:02; 40:01; 44:02; |
| | YKHWPQIAQF | 24:02 |
| | GLPNNTASWF | 24:02 |
| | NKHIDAYKTF | 24:02 |
| | PNFKDQVILL | 24:02 |
| | QGLPNNTASW | 24:02 |
| | SSPDDQIGYY | 01:01 |
| | VTPSGTWLTY | 01:01 |
| | GTGPEAGLPY | 01:01; 11:01 |
| | NSSPDDQIGY | 01:01 |
| | QGTTLPKGFY | 01:01 |
| | DLSPRWYFYY | 01:01 |
| | ILLNKHIDAY | 01:01 |
| | KTFPPTEPKK | 03:01; 11:01 |
| | YKTFPPTEPK | 03:01; 11:01 |
| | ATEGALNTPK | 03:01; 11:01 |
| | KLDDKDPNFK | 03:01; 11:01 |
| | LLNKHIDAYK | 03:01 |
| | RIRGGDGKMK | 03:01 |
| | KMKDLSPRWY | 03:01 |
| | KFPRGQGVPI | 07:02 |
| | NPANNAAIVL | 07:02; 08:01 |
| | APSASAFFGM | 07:02 |
| | RPQGLPNNTA | 07:02 |
| | RNPANNAAIV | 07:02 |
| | FPRGQGVPIN | 07:02 |
| | RGPEQTQGNF | 07:02 |
| | SASAFFGMSR | 11:01 |
| | KKSAAEASKK | 11:01 |
| | RSKQRRPQGL | 08:01 |
| | DPNFKDQVIL | 08:01 |
| | DLKFPRGQGV | 08:01 |
| | NQRNAPRITF | 08:01; 40:01 |
| | DFSKQLQQSM | 08:01 |
| | MEVTPSGTWL | 40:01; 44:02 |
| | GMEVTPSGTW | 40:01; 44:02 |
| | GPEAGLPYGA | 40:01; 44:02 |
| | GDAALALLLL | 40:01 |

**b**

| | Peptide | HLA-DR |
|---|---|---|
| RBD | RVVVLSFELLHAPATVCGP | 01:01 |
| | FNGLTGTGV | 01:01 |
| | IRGDEVRQI | 03:01 |
| | LKPFERDISTEIYQA | 03:02 |
| | FQPTNGVGY | 07:01 |
| Nucleoprotein | KDGIIWVATEGALNTPKD | 01:01 |
| | PANNAAIVLQLPQGTTLPKGF | 01:02 |
| | VTLLPAADL | 01:01; 01:02 |
| | DDQIGYYRRATRRIRGGD | 01:03 |
| | FYAEGSRGG | 01:03 |
| | LTYTGAIKLDDKDPNFKDQ | 03:01 |
| | DAALALLLLDRLNQLESKMS | 03:02; 04:11 |
| | TPSGTWLTYTGAIKLDDKDPN | 07:01 |
| | YNVTQAFGR | 07:01 |
| | FAPSASAFF | 07:01 |

**c**

| | Peptide | MHC-I |
|---|---|---|
| RBD | IVRFPNITNL | H-2K$^b$; H-2D$^b$ |
| | VGYQPYRVVV | H-2K$^b$ |
| | YSVLYNSASF | H-2D$^b$ |
| | STPCNGVEGF | H-2D$^b$ |
| | SKVGGNYNYL | H-2D$^b$ |
| Nucleo-protein | LSPRWYFYYL | H-2K$^b$ |
| | AQFAPSASAF | H-2K$^b$ |
| | ALALLLLDRL | H-2D$^b$ |
| | TRNPANNAAI | H-2D$^b$ |
| | VLQLPQGTTL | H-2D$^b$ |

**d**

| | Peptide | MHC-II |
|---|---|---|
| RBD | TNVYADSFVIRGDEV | H-2IA$^b$ |
| | PDDFTGCVIAWNSNN | H-2IA$^b$ |
| | FELLHAPATVCGPKK | H-2IA$^b$ |
| Nucleo-protein | YFYYLGTGPEAGLPY | H-2IA$^b$ |
| | TASWFTALTQHGKED | H-2IA$^b$ |
| | RITFGGPSDSTGSNQ | H-2IA$^b$ |
| | GIIWVATEGALNTPK | H-2IA$^b$ |
| | KEDLKFPRGQGVPIN | H-2IA$^b$ |
| | EGALNTPKDHIGTRN | H-2IA$^b$ |

**Extended Data Figure 1.** Amino acid sequences of putative epitopes from RBD and N proteins found by *in silico* epitope prediction. The peptides are listed according to specificity for (A) HLA-ABC, (B) HLA-DR, (C) murine MHC-I and (D) MHC-II alleles.

**Extended Data Fig. 2**

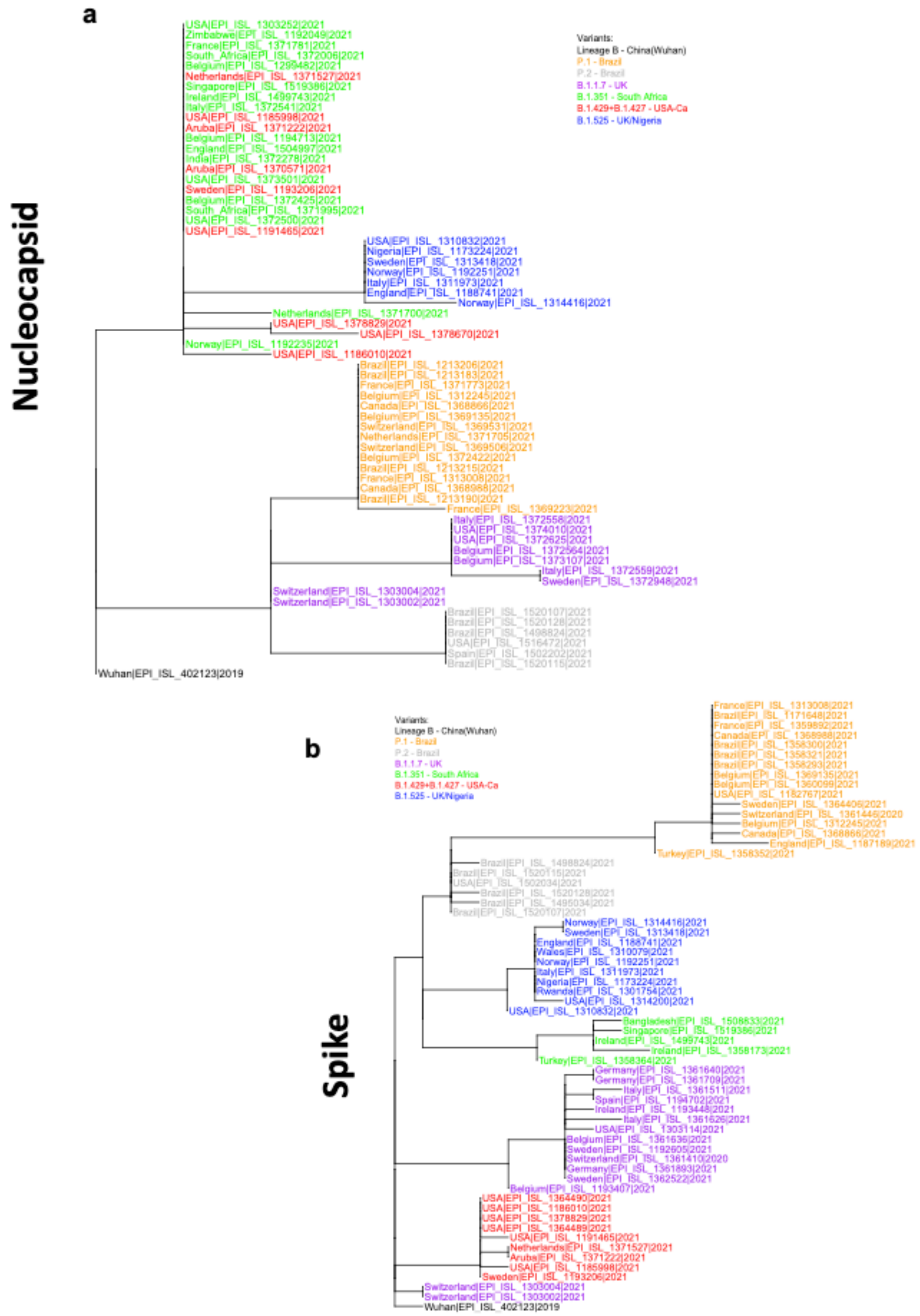


**Extended Data Figure 2. a,b,** The dendrograms were constructed based on nucleocapsid **(a)** and spike **(b)** nucleotide sequences from 61 and 63 SARS-CoV-2 isolates found in different geographic areas worldwide. These sequences belong to the five most relevant groups of variants of SARS-CoV-2 presented in GISAID as UK - VUI202012 / 01 GRY-B.1.1.7, South Africa - GH/501Y.v2-B.1.351, Brazil - GR/501Y.V3-P.1, USA-Ca - GH/452R. V1-B.1.429 + B.1.427, UK/Nigeria - G/484K.V3-B.1.525 and the variant P.2 from Brazil. The lineage B was defined as "outgroup". The predicted amino acid sequence of the nucleocapsid and spike proteins from same isolates were used to build the Needle plot shown in **Figure 1a**.

Extended Data Fig. 3

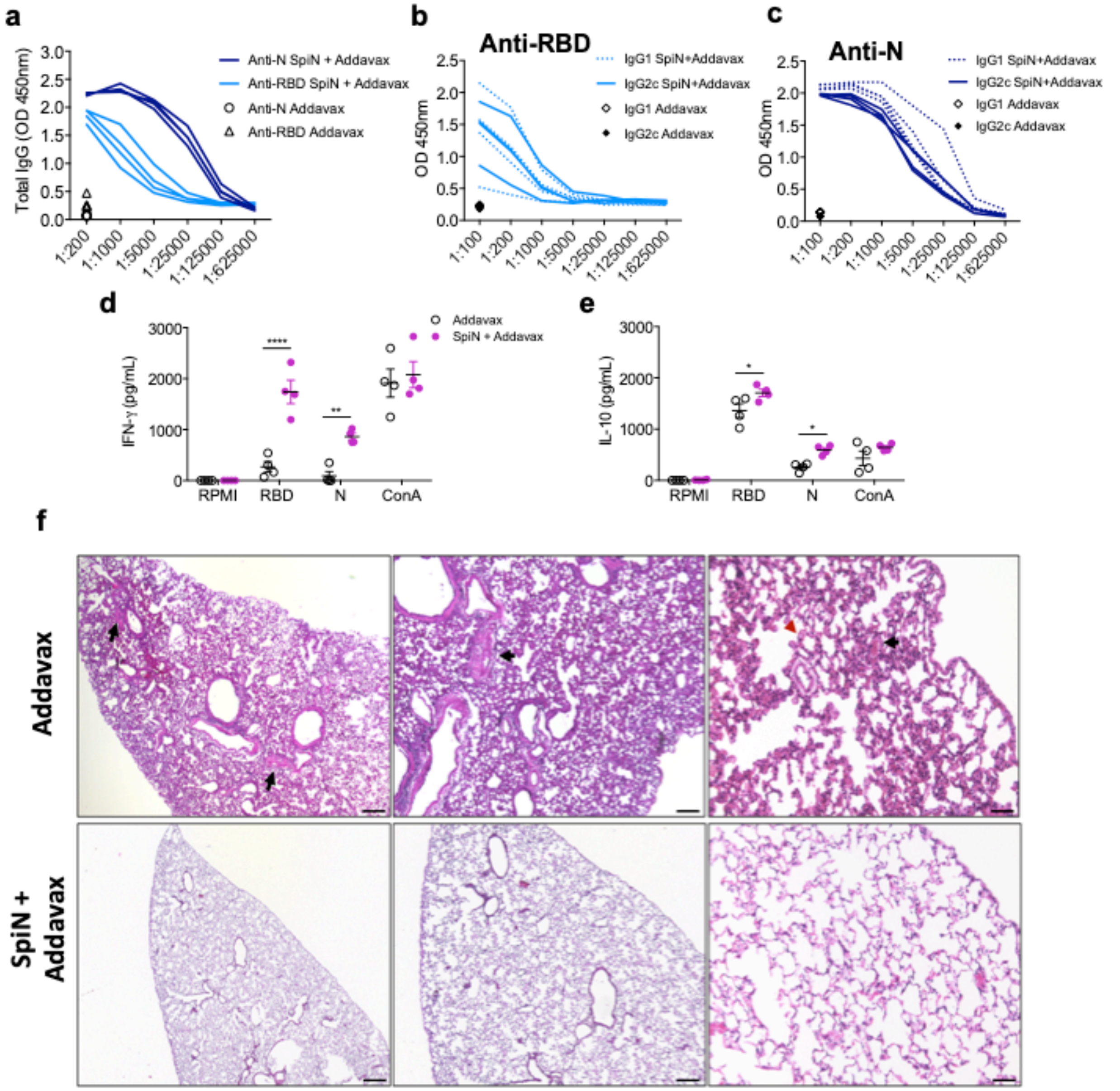


**Extended Data Figure 3. Evaluation of immune response and protection in mice immunized with SpiN associated with Addavax. a,** Total IgG antibodies anti-N and RBD measured in the sera from immunized mice. The production of IgG1 and IgG2c specific for RBD **(b)** and N **(c)** proteins was evaluated in the sera from mice vaccinated with SpiN + Addavax. The levels of IFNγ **(d)** and IL-10 **(e)** produced by splenocytes after stimulation with RBD or N proteins. **f,** Histopathological sections of the lungs from mice challenged with the Wuhan strain of SARS-CoV-2. Animals were administered with Addavax only (upper panel) or with SpiN + Addavax (bottom panel). * P < 0.05, ** P < 0.01 and *** P < 0.001.
